# Genome-wide meta-analyses of stratified depression in Generation Scotland and UK Biobank

**DOI:** 10.1101/130229

**Authors:** Lynsey S. Hall, Mark J. Adams, Aleix Arnau-Soler, Toni-Kim Clarke, David M. Howard, Yanni Zeng, Gail Davies, Saskia P. Hagenaars, Ana Maria Fernandez-Pujals, Jude Gibson, Eleanor M. Wigmore, Thibaud S. Boutin, Caroline Hayward, Generation Scotland, Major Depressive Disorder Working Group of the Psychiatric Genomics Consortium, David J. Porteous, Ian J. Deary, Pippa A. Thomson, Chris S. Haley, Andrew M. McIntosh

**Author notes:** Correspondence to: Dr. Lynsey S. Hall, Division of Psychiatry, University of Edinburgh, Kennedy Tower, Royal Edinburgh Hospital, Edinburgh, EH10 5HF, UK.

## Abstract

Few replicable genetic associations for Major Depressive Disorder (MDD) have been identified. However recent studies of depression have identified common risk variants by using either a broader phenotype definition in very large samples, or by reducing the phenotypic and ancestral heterogeneity of MDD cases. Here, a range of genetic analyses were applied to data from two large British cohorts, Generation Scotland and UK Biobank, to ascertain whether it is more informative to maximize the sample size by using data from all available cases and controls, or to use a refined subset of the data - stratifying by MDD recurrence or sex. Meta-analysis of GWAS data in males from these two studies yielded one genome-wide significant locus on 3p22.3. Three associated genes within this region (*CRTAP, GLB1*, and *TMPPE*) were significantly associated in subsequent gene-based tests. Meta-analyzed MDD, recurrent MDD and female MDD were each genetically correlated with 6 of 200 health-correlated traits, namely neuroticism, depressive symptoms, subjective well-being, MDD, a cross-disorder phenotype and Bipolar Disorder. Meta-analyzed male MDD showed no statistically significant correlations with these traits after correction for multiple testing. Whilst stratified GWAS analysis revealed a genome-wide significant locus for male MDD, the lack of independent replication, the equivalent SNP-based heritability estimates and the consistent pattern of genetic correlation with other health-related traits suggests that phenotypic stratification in currently available sample sizes is currently weakly justified. Based upon existing studies and our findings, the strategy of maximizing sample sizes is likely to provide the greater gain.

## Introduction

Major Depressive Disorder (MDD) is a frequently disabling, chronic disorder for which there is substantial evidence of a genetic contribution to its liability^1^. Until recently, the largest international mega-analysis of clinically diagnosed MDD (9 240 MDD cases and 9 519 controls) yielded no genome-wide significant findings^2^. Given the success of similarly sized studies for other adult psychiatric disorders^3,4^, this study suggested that MDD is an extensively heterogeneous phenotype. This heterogeneity, in addition to the relatively high prevalence and low heritability of MDD, impacts substantially on the statistical power to detect genetic effects^1,5^. Possible means of improving statistical power include stratifying the phenotype into potentially more homogeneous subtypes, or considerably increasing the sample size whilst accepting a broader phenotype. Both of these approaches have since identified associations between genetic variants and MDD^6-9^.

A genome-wide association study (GWAS) from the CONVERGE consortium identified two genome-wide significant loci (chromosome 10q21.3 and 10q26.13) using a severe depressive phenotype (5 053 cases and 5 337 controls) in female Han Chinese individuals, treated in a hospital setting^6^. A subsequent study by the Psychiatric Genomics Consortium (PGC) stratified the PGC MDD mega-analysis sample^2^ by age of onset and identified a risk conferring locus at chromosome 3q27.2 in individuals with onset after 27 years of age^9^.

In contrast to the above study designs, two studies have utilized larger sample sizes with less detailed structured clinical assessments. The first of these studies came from the CHARGE consortium and employed a quantitative assessment of depressive symptoms using the Center for Epidemiological Studies Depression Scale. In a combined dataset of 51 258 individuals, a genome-wide significant locus was identified at chromosome 5q21.2^7^. More recently, Hyde et al^8^ conducted a GWAS using 23andMe data of self-reported depression in 45 773 cases and 106 354 controls, revealing 15 genome-wide significant loci. The genetic correlation (rG) between the 23andMe depression phenotype and the clinical phenotype reported by the PGC was rG(SE)=0.73(0.09), suggesting a strong association between the additive genetic components of each trait. Previous work comparing self-reported depression and clinically defined MDD in Generation Scotland: Scottish Family Health Study (GS:SFHS) also provides evidence that these traits have substantially overlapping common genetic architectures^10^.

The findings from these four studies support both phenotypic stratification and increased sample size as strategies which may help reveal the underlying architecture of MDD.

The international collaborative efforts by groups such as the PGC and the development of large-scale biobanks with genetic and extensive phenotypic information will ensure ever increasing sample numbers. It is therefore timely to investigate the contrasting strategies that may be employed in the analysis of these emerging datasets. This is particularly relevant to MDD, given the disparate strategies that have recently shown success in identifying genome-wide significant loci^6,8,9^. In the current study, we sought to compare these strategies by conducting GWAS of depression and stratified subtypes in two UK-based cohorts: GS:SFHS^11,12^ and UK Biobank (UKB)^13,14^. To maximize the sample size, an unstratified analysis was initially conducted. This used MDD diagnostic information obtained at structured clinical interview^15^ in GS:SFHS (2 603 cases, 16 122 controls), and a probable MDD phenotype obtained from a touchscreen questionnaire^16^, previously validated by Smith et al^17^, in UKB (8 248 cases, 16 089 controls). Subsequent analyses stratified the phenotype on the basis of recurrence or sex. Each approach was evaluated by performing genome-wide meta-analyses, and comparing the successful identification of any genome-wide variants, the SNP-based heritability, the genetic correlation with other traits using LD score regression, and the variance explained by polygenic profile scores for MDD derived from three independent cohorts for each MDD phenotype subgroup.

## Materials and Methods

This study analyzed data from Generation Scotland: The Scottish Family Health Study (GS:SFHS), (http://www.generationscotland.co.uk) and UK Biobank (UKB), (http://www.ukbiobank.ac.uk). GS:SFHS received ethical approval from the NHS Tayside Committee on Medical Research Ethics (REC Reference Number: 05/S1401/89). UK Biobank received ethical approval from the Research Ethics Committee (REC Reference Number: 11/NW/0382). The present analyses were conducted under UK Biobank data application number 4844.

## Participants

### Generation Scotland: The Scottish Family Health Study (GS:SFHS)

GS:SFHS is a family- and population-based study consisting of 23 690 participants recruited via general medical practices across Scotland. The recruitment protocol and sample characteristics are described in detail elsewhere^11,12^. Briefly, participants were over 18 years old, and not ascertained on the basis of having any particular disorder. A diagnosis of depression (MDD) was made using the structured clinical interview for DSM-IV disorders (SCID)^15^. Participants who answered yes to either of the two screening questions were invited to continue the interview, which provided information on the presence or absence of a lifetime history of MDD, age of onset and number of depressive episodes. Participants who answered no to both screening questions or who completed the SCID but did not meet the criteria for depression were assigned control status. Case definition was further refined through NHS data linkage. Controls with a history of antidepressants or who had been referred to a secondary psychiatric care centre (n=1072) were excluded, as were cases who had received a previous diagnosis of schizophrenia or bipolar disorder (n=47). This resulted in 2 603 depression cases (of which 1 289 were recurrent) and 16 122 controls. Stratification by sex resulted in 1 859 female cases, 9 159 female controls, 770 male cases and 6 958 male controls.

### UK Biobank (UKB)

UK Biobank^13^ is a population-based health research resource consisting of approximately 500 000 people, aged between 40 and 69 years, who were recruited between the years 2006 and 2010 from across the UK^14^. Of these, 152 729 individuals were included in the first genotype data release. In the current study we restricted the sample to individuals of white British ancestry. Participants who were also in GS:SFHS, their relatives and relatives of remaining UKB participants (relatives: up to and including 3rd degree) were identified by a kinship coefficient ≥0.0442, using the KING toolset^18^, and subsequently excluded (n=7 698). Depression case/control status was assessed in 172 751 of the 500 000 individuals using a self-diagnosed touchscreen questionnaire. Case status was defined as either ‘probable single lifetime episode of major depression’ or ‘probable recurrent major depression (moderate and severe)’. Control status was defined as ‘no mood disorder’, as described by Smith *et al*^17^. 149 847 individuals had sufficient data to allow an assessment of case/control status. Individuals with probable bipolar disorder (n=1 615) or mild depressive/manic symptoms (n=26 847) were excluded. After exclusions outlined above, this resulted in 8 248 depression cases (of which 6 056 were recurrent) and 16 089 controls. Stratification by sex resulted in 5 138 female cases, 7 013 female controls, 3 110 male cases and 9 076 male controls. Further information on sample collection, genotyping and assessment of the depression phenotype in GS:SFHS and UKB are provided in the Supplementary Methods.

## Imputation and quality control

### GS:SFHS

Autosomal genotype data were available for all GS:SFHS individuals in the present study (n=18 725). Genotypes were imputed using the Haplotype Reference Consortium reference panel (HRC.r1-1)^19^ via the Sanger Imputation Server pipeline (https://imputation.sanger.ac.uk). Prior to imputation, individuals with missingness ≥3% were excluded, as were SNPs with a call rate of ≤98%, Hardy Weinberg Equilibrium (HWE) P-value ≤1 × 10^−6^, and a minor allele frequency (MAF) ≤1%. Phasing of genotype data was performed using the SHAPEIT2 algorithm^20^ utilizing the duoHMM option, which refines phasing by utilizing pedigree information. Imputation was performed using PBWT software^21^. Multi-allelic variants, monomorphic variants and SNPs with an imputation INFO score <0.8 were removed^22^. Population outliers (more than 6SDs from the mean of the first principal component (PC)) were identified and removed from the sample^23^, as were one from each of 52 monozygotic twin pairs, identified by IBD (preferentially retaining cases), and 7 individuals who matched samples from the Psychiatric Genomics Consortium, identified using genotype checksums^24^. After imputation, individuals with missingness ≥2%, and genotype with a call rate of ≤98%, MAF ≤0.5% and HWE P-value ≤1E-05 were excluded using PLINK version 1.9.^25,26^. Strand ambiguous SNPs with 40%≤MAF≤50% were also excluded.

### UKB

Autosomal genotypes were available for all UKB individuals in the present study (n=24 337). Pre-imputation QC, phasing and imputation are described elsewhere^27^. In brief, prior to phasing, multiallelic SNPs or those with MAF ≤1% were removed. Phasing of genotype data was performed using a modified version of the SHAPEIT2 algorithm^28^. Imputation to a reference set combining the UK10K haplotype and 1000 Genomes Phase 3 reference panels^29^ was performed using IMPUTE2 algorithms^30,31^. A further QC protocol was then applied at the Wellcome Trust Centre for Human Genetics before the data was released, as described elsewhere^32^. The analyses presented here were restricted to autosomal variants with an imputation INFO score ≥ 0.9 and MAF ≥0.5%.

Of the SNPs which passed QC in each dataset, only SNPs in common between both datasets were used in subsequent analyses, with allele and strand in GS:SFHS harmonized to be consistent with UKB, resulting in 7 105 178 autosomal SNPs.

### Statistical analysis

All analyses were performed on four subsets of the data: all available cases and controls (MDD), all controls and recurrent cases (rMDD), female controls and cases (fMDD), and male controls and cases (mMDD). The total sample size for each depression subgroup was n=43 062 for MDD, n=39 556 for rMDD, n=23 169 for fMDD and n=19 886 for mMDD. The number of cases and controls and demographic information for these subsets are shown in Supplementary Table 1.

## Association analysis

### GS:SFHS

Genome-wide association analysis (GWAS) of MDD, rMDD, fMDD and mMDD in GS:SFHS were conducted using mixed linear model based association (MLMA) analysis^33^, implemented in GCTA (v1.25.)^34^. To account for population structure, two genomic relationship matrices (GRMs) were used, as this method allows the inclusion of closely and distantly related individuals in genetic analyses^35^. The first GRM included pairwise relationship coefficients for all individuals. The second GRM had off-diagonal elements < 0.05 set to 0. GRMs were created using the mixed linear model with candidate marker excluded (MLMe) approach, where GRMs are calculated excluding SNPs located on the chromosome under analysis^33^. No fixed effects covariates were fitted in this analysis as sex was being assessed as a stratifier, and the two GRMs adequately accounted for population stratification (tested using univariate LD Score Regression^36^). MLMA employs restricted maximum likelihood methods on the linear scale. As such, test statistics (betas and their corresponding standard errors) were transformed to Odds Ratios and their corresponding 95% Confidence Intervals on the liability scale using a Taylor transformation expansion series^37,38^. Further details of GWAS can be found in the Supplementary Methods.

### UKB

GWAS of MDD, rMDD, fMDD and mMDD in UKB were conducted using logistic regression, implemented in PLINK v1.9^25^. Assessment centre, genotype array and batch were fitted as fixed effects. The first 8 PCs (out of 15) supplied by UKB were also fitted, as visual inspection indicated that these PCs resulted in multiple clusters, indicating structure in the data.

### Meta-analysis, variant look-up and gene-based analysis

The meta-analysis of GS:SFHS and UKB was conducted using the classical inverse-variance approach, which weights effect sizes by sampling distribution, implemented in the METAL package^39^. SNPs with a meta-analysis P-value of P≤1E-05 were subjected to clump-based linkage disequilibrium pruning using PLINK^25^ using an LD r^2^ cut off of 0.1 and a 500kb sliding window to create SNP sets of approximately independent “lead SNPs”. All SNPs which surpassed genome-wide significance were entered into the NHGRI-EBI catalog of published GWAS^40,41^ (www.ebi.ac.uk/gwas/) to observe whether these SNPs had been previously observed in association analysis.

Gene-based analysis was performed for MDD, rMDD, fMDD, and mMDD using MAGMA^42^. The gene-based statistics were derived using the summary statistics from each meta-analysis. Genetic variants were assigned to genes based on their position according to the NCBI 37.3 build, with a gene boundary defined by an extended region between 20 kb upstream of transcript start site and 20kb downstream of transcript end site for each of the genes. This resulted in a total of 18 111 genes for MDD, fMDD, and mMDD, and 17 225 genes for rMDD being analyzed. The European panel of the 1000 Genomes data (phase 1, release 3) was used as a reference panel to account for linkage disequilibrium^43^. A genome-wide significance threshold for gene-based associations was calculated using the Bonferroni method (α=0.05/18 111; P<2.76 × 10^−6^ for MDD, fMDD and mMDD; α=0.05/17 225; P<2.90 × 10^−6^ for rMDD).

### Pathway and functional genomic analyses

Pathway and functional genomic analyses were performed using the GWAS results for each of the MDD meta-analyses. These included DEPICT analyses^44^, reference to RegulomeDB^45^ (http://www.regulomedb.org/) and to the Genotype-Tissue Expression Portal (http://www.gtexportal.org) for independent SNPs with P<1.0 × 10^−5^ and all genome-wide significant SNPs (P<5.0 × 10^−8^, nSNPs = 6). Further information on pathway and functional genomic analysis can be found in the Supplementary Methods.

### Heritability, polygenicity and cross-trait genetic correlations

Univariate GCTA-GREML^46^ analyses were used to estimate the proportion of variance explained by all common (MAF>1%) SNPs for each of the depression phenotypes. A relatedness cutoff of 0.05 was used in the generation of the genetic relationship matrix, as including close relatives inflates heritability estimates^47^. This did not alter the sample size in UKB due to previous sample filtering, however in GS:SFHS this reduced the sample size by 38.5-58.4% (Supplementary Table 15). In GS:SFHS, the first 20 PCs were fitted as fixed effects. In UKB, batch, recruitment centre and the first 8 PCs were fitted. Univariate Linkage Disequilibrium Score regression (LDSR), implemented in LD Score (v1.0.0.)^36^, was applied to GWAS summary statistics to evaluate the proportion of inflation in the test statistics caused by confounding biases, such as population stratification, relative to genuine polygenicity. This method also provides an estimate of SNP-based heritability. Pre-computed LD Scores were used, estimated from the European-ancestry samples in the 1000 Genomes Project^43^. To obtain heritability estimates on the liability scale, sample and population prevalence estimates were used. Sample prevalence estimates were calculated as the proportion of cases in each subset. Population prevalence estimates were derived from the literature^48-50^. Prevalence estimates used in GCTA-GREML and LDSC are given in Supplementary Table 1. Genetic correlations between meta-analyzed depression subgroups and 200 health-related traits were calculated using bivariate LDSR^51^, implemented in the LD Hub software^52^. Traits derived from non-Caucasian or mixed ethnicity samples were removed prior to analysis. False discovery rate (FDR) correction was applied across the 800 tests to correct for multiple testing^53^.

### Polygenic profiling analysis

To test the association of MDD-associated alleles with each subtype of MDD in GS:SFHS and UKB, summary statistics for major depressive disorder from the Psychiatric Genomics Consortium (unpublished), UKB and GS:SFHS (from the current study) were used to provide weights for polygenic profile scores (PGS).

PGS for GS:SFHS and UKB individuals were derived at 5 GWAS P-value thresholds (P_T_<0.01, <0.05, <0.1, <0.5 and all SNPs) using PRSice^54^. Genotyped SNPs (with MAF >1%) were subjected to clump-based linkage disequilibrium (LD) pruning, using an LD r^2^ cut off of 0.25 and 200kb sliding window to create SNP sets in approximate linkage equilibrium. PGS were then standardized to have a mean of zero and a unit standard deviation.

In GS:SFHS, the associations between PGS and MDD, rMDD, fMDD and mMDD were tested using a mixed linear model, covarying for the first 20 PCs to account for population stratification. Prior to this analysis, the requisite number of PCs was established using a stepwise linear regression approach, adding one PC at a time, and using a likelihood-ratio test (LRT), the output of which was assessed against a mixed 0.5(χ^2^)+0.5(0) distribution^55^. An additive genetic component was fitted as a random effect to account for the increased relatedness within GS:SFHS. To ensure that common environment was adequately modeled, models incorporating shared parent-offspring, sibling, and spousal environmental components as additional random effects were tested using a stepwise LRT approach, however no environmental component improved model fit. Further details of mixed linear model selection are provided in the Supplementary Methods. F-statistics, degrees of freedom, effect sizes, Z-scores and P-values were derived using the Wald Conditional F-test^56^, in ASReml-R^57^.

In UKB, the association between PGS and MDD, rMDD, fMDD and mMDD was tested in a generalized linear model framework by regressing the PGS onto the phenotype, covarying for assessment centre, genotype array and batch and the first 8 PCs.

FDR correction was applied across the 80 tests to correct for multiple testing^53^. For both GS:SFHS and UKB, trait variance explained by the PGS was calculated using: (var(*x* × *β*))/var(*y*), where *x* was the standardized PGS, *β* was the corresponding regression coefficient and *y* was the phenotype^58^.

## Results

### Meta-analysis of depression in GS:SFHS and UKB

One genomic region on chromosome 3p22.3 achieved genome-wide significant in the males only case/control (mMDD) analysis (index SNP rs4478037, β(SE)=0.29(0.05), P=2.29 × 10^−8^). No genome-wide significant SNPs associated with any phenotype in currently published GWAS available via the NHGRI-EBI catalog. One variant (rs7613051) within the local genomic region (defined 3:33000000-33200000) has previously shown an association with Atopic dermatitis^59^, however this SNP is not in LD with the genome-wide significant SNPs (r^2^<0.1). Manhattan plots are shown for each trait in Figure 1, and summary statistics for independent loci with a meta-analysis association P≤1 × 10^−6^ are shown in Table 1. A regional association plot for genome-wide significant index SNP, rs4478037, is shown in Figure 2. Regional association plots for this SNP in other depression subtypes are shown in Supplementary Figure 4, demonstrating that this locus does not replicate in any other depression subtype (minimum P=8.05 × 10^−4^ in MDD, β(SE)=0.10(0.05)). Full details of all independent loci used in downstream analyses (P≤1 × 10^−5^) are shown in Supplementary Tables 4-11. The QQ plots (Supplementary Figure 3) demonstrate λ_GC_ ranged from 1.02 - 1.06, comparable to the value (1.056) observed in the PGC mega-analysis of MDD^2^. Univariate LDSR analyses estimated that meta-analyzed MDD subtypes had mean chi-squared statistic (μχ^2^) values ranging from 1.018 (mMDD) to 1.062 (MDD) with a Ratio, defined as (Intercept - 1)/(μχ^2^-1), ≤ 0.35 across subtypes, indicating that any inflation in μχ^2^ can be attributed to polygenicity rather than residual population stratification^36^.

**Figure 1.**
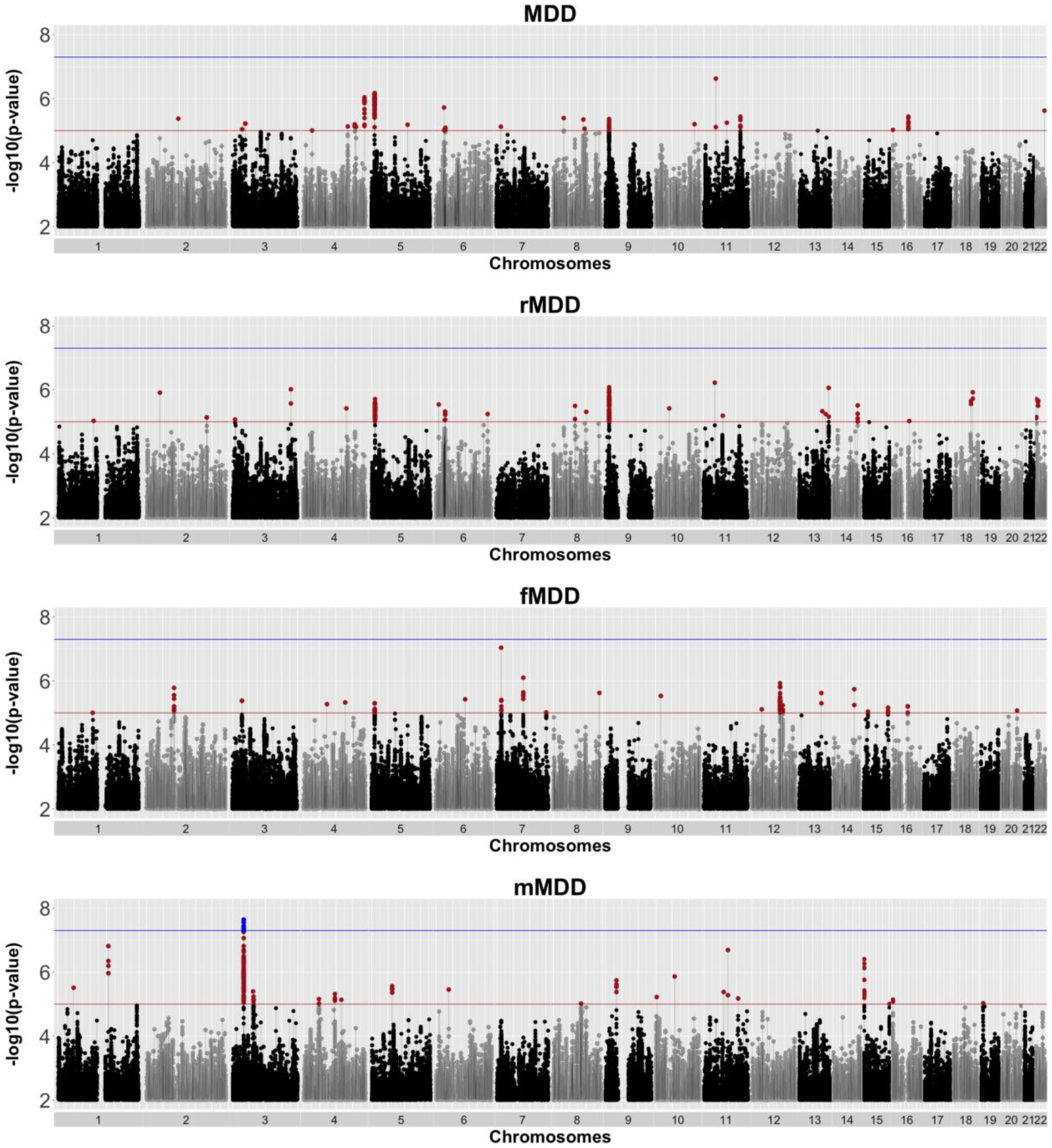
Manhattan plot of P-values from SNP-based association meta-analysis of all depression cases and controls (MDD, n=43 062), recurrent only cases and all controls (rMDD, n=39 556), females only cases and controls (fMDD, n=23 169) and males only cases and controls (mMDD, n=19 886). The blue line indicates the threshold for genome-wide significance (P<5 × 10^−8^), the red line indicates the threshold for suggestive significance (P<1 × 10^−5^).

**Figure 2.**
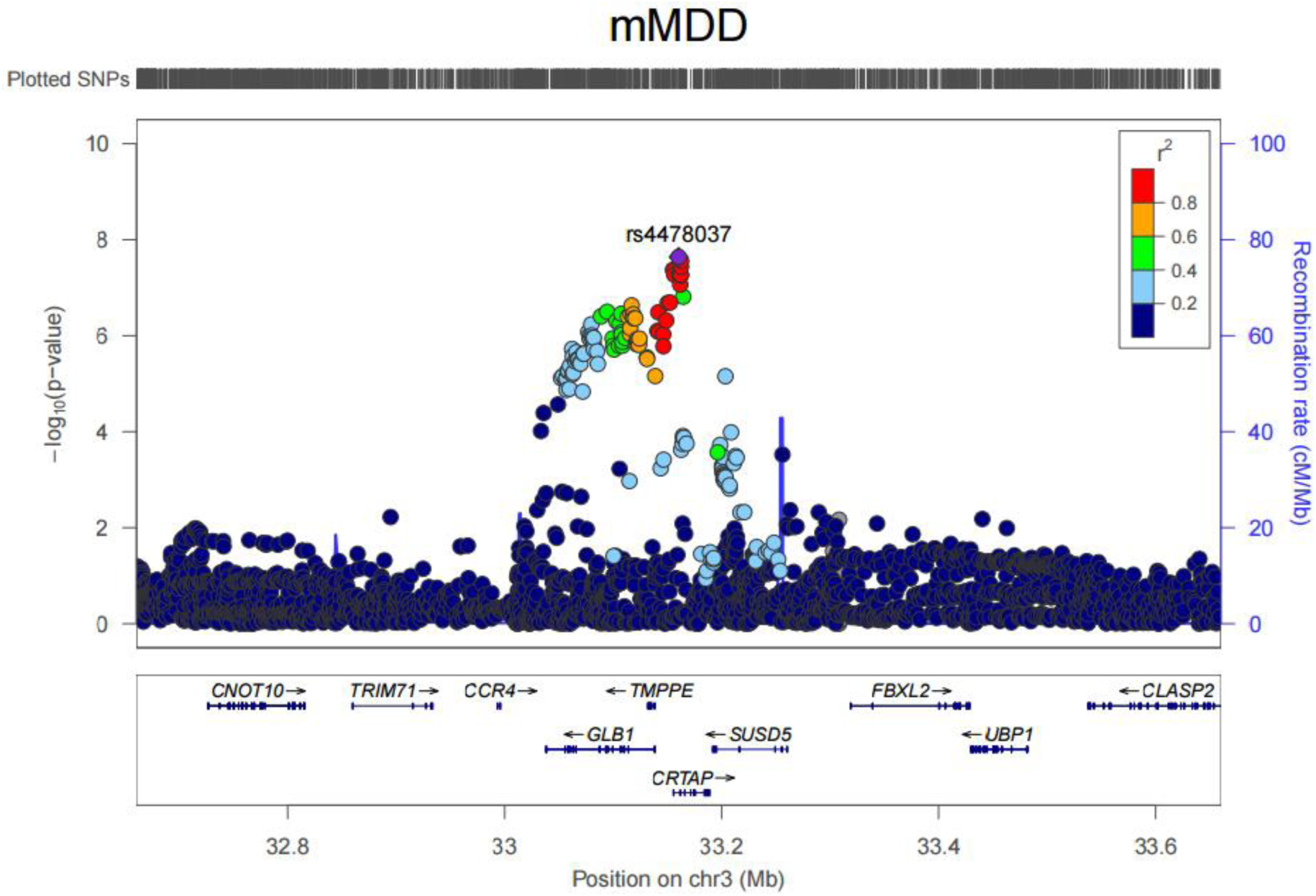
Regional association plot for rs4478037, an intronic SNP in *CRTAP*, and the top ranking SNP (rs4478037, P=2.37 × 10^−8^) in GWAS of depression in males only.

**Table 1.** Summary statistics for SNPs with association P-value ≤1 × 10^−6^ for depression (MDD), recurrent depression (rMDD), depression in females only (fMDD) and depression in males only (mMDD), sorted within phenotype by genomic positions according to UCSC hg19/NCBI Build 37. Column A1/A2 contains the reference and alternate alleles for the index SNP, respectively. The meta-analysis minor allele frequency (Freq) and regression coefficient (β) columns pertain to the reference allele (A1). Chr and Position denote the location of the index SNP. SE is the standard error for β. The direction of effect of the index SNP in GS:SFHS and UKB is shown in the Direction column. The final column, Genes, indicates protein-coding reference sequence genes within 10kb of the associated loci.

| Trait | SNP | CHR | POS | A1/A2 | Freq | $\beta$(se) | P | Direction | Genes |
| --- | --- | --- | --- | --- | --- | --- | --- | --- | --- |
| MDD | rs56390503 | 4 | 187552576 | T/C | 0.11 | 0.13 (0.03) | 9.08E-07 | ++ | *FAT1* |
|  | rs2964802 | 5 | 10820843 | T/C | 0.28 | 0.09 (0.02) | 6.73E-07 | ++ | - |
|  | rs11033303 | 11 | 35871266 | A/G | 0.37 | 0.09 (0.02) | 2.37E-07 | ++ | - |
| rMDD | rs2291479 | 3 | 178174944 | A/C | 0.4 | -0.10 (0.02) | 9.65E-07 | -- | - |
|  | rs10959631 | 9 | 11220986 | T/C | 0.2 | -0.12 (0.02) | 8.34E-07 | -- | - |
|  | rs11033303 | 11 | 35871266 | A/G | 0.37 | 0.11 (0.02) | 6.02E-07 | ++ | - |
|  | rs4438172 | 13 | 111448658 | A/T | 0.25 | 0.11 (0.02) | 8.72E-07 | ++ | - |
| fMDD | rs9648182 | 7 | 13794849 | A/T | 0.12 | -0.17 (0.03) | 9.14E-08 | -- | - |
|  | rs17176546 | 7 | 81880914 | A/G | 0.04 | 0.26 (0.05) | 7.93E-07 | ++ | *CACNA2D1* |
| mMDD | rs115736167 | 1 | 155266609 | C/G | 0.02 | -0.46 (0.09) | 1.54E-07 | -- | *PKLR* |
|  | rs4478037 | 3 | 33160407 | A/G | 0.08 | 0.29 (0.05) | 2.29E-08 | ++ | *CRTAP* |
|  | rs113485090 | 11 | 73572495 | A/G | 0.04 | -0.32 (0.06) | 2.05E-07 | -- | *MRPL48* |
|  | rs1380551 | 15 | 24124704 | A/G | 0.15 | -0.18 (0.04) | 3.96E-07 | -- | - |

### Gene based analysis of MDD subtypes

Three genes at chromosome 3p22.3 (*CRTAP, GLB1* and *TMPPE*) were significantly associated with mMDD after Bonferroni correction (Supplementary Table 12). There were no significant gene-based associations with MDD, rMDD or fMDD.

### Pathway and functional genomic analyses

Gene set enrichment analysis of SNPs with meta-analysis P≤1 × 10^−5^, as implemented in DEPICT, indicated a role for two gene sets at FDR<0.05 in mMDD (Supplementary Table 13): GO terms carboxylic acid binding and CNTN1 PPI subnetwork. No other significant results were observed for tissue enrichment or gene prioritization.

Using the GTEx database (http://www.broadinstitute.org/gtex/), 25 multi-tissue cis-eQTL associations were identified for 16 independent lead SNPs with meta-analysis P<1 × 10^−5^ (Supplementary Tables 4, 6, 8, 10, and 14). The 5 genome-wide significant SNPs identified for mMDD show eQTL evidence for the genes *GLB1* and *CRTAP*. Random effect meta-analysis of multi-tissues for the most significant mMDD SNP, rs447803, yielded P=3.58 × 10^−9^ and 8.45 × 10^−29^ for *GLB1* and *CRTAP* respectively. For this study, data mining of regulatory elements was restricted to normal cell lines/tissues. There was evidence of regulatory elements (Regulome DB score<4) for 6 of the lead SNPs with meta-analysis P≤1 × 10^−5^ (MDD: rs10736455, rs73249855, rs8050755; rMDD: rs60716536; fMDD: rs11613048; mMDD: rs74002781). Of the 6 SNPs which achieved genome-wide significance in the meta-analysis of mMDD, 2 SNPs (rs11558338 and rs6809511) showed evidence of transcription factor binding, position weight matrix, histone modification, DNase hypersensitivity, and FAIRE regulatory elements. Evidence of regulatory evidence for all independent SNPs with meta-analysis P≤1 × 10^−5^ are shown in Supplementary Table 14.

### Estimating SNP-based heritability and polygenicity

Using GCTA-GREML methods^46^, the SNP-based heritability (h^2^_SNP_) estimates in UKB were consistent and significant across MDD subtypes, with h^2^_SNP_ (SE) estimates of MDD = 0.20(0.04); rMDD = 0.20(0.03); fMDD = 0.22(0.06) and mMDD = 0.18(0.06). Due to the unrelated subset of individuals in GS:SFHS being markedly smaller than the full sample (n_max_=7 795), the heritability estimates were non-significant. Results from GCTA-GREML are shown for MDD subtypes in Supplementary Table 15. LDSR yielded lower h^2^_SNP_ estimates than GCTA-GREML methods (Supplementary Table 16). This is to be expected as LDSR utilizes summary scores, which have usually been subjected to genomic control, as opposed to full SNP data.

### Genetic correlation with health-related traits

Bivariate LDSR showed nominally significant (P<0.05) genetic correlations (rG) between meta-analyzed MDD and 28 of the 200 health-related traits assessed. Of these, 8 traits survived multiple testing correction: neuroticism (rG(SE)=0.67(0.07); P=7.06 × 10^−21^), depressive symptoms (rG(SE)=0.81(0.09); P=1.72 × 10^−19^), subjective wellbeing (rG(SE)=-0.56(0.08); P=9.12 × 10^−13^), age at first birth (rG(SE)=-0.35(0.05); P=1.92 × 10^−10^), major depressive disorder (rG(SE)=0.67(0.12); P=4.57 × 10^−8^), PGC cross-disorder analysis (rG(SE)=0.46(0.09); P=8.60 × 10^−8^), bipolar disorder (rG(SE)=0.32(0.08); P=4.35 × 10^−5^) and systemic lupus erythematosus (rG(SE)=0.28(0.08); P=8.00 × 10^−4^). The majority of these traits (with the exception of age at first birth and systemic lupus erythematosus) also had similar rG and significantly associated with recurrent and female MDD, but not male MDD (Supplementary Table 17). Figure 3 shows the genetic correlation of meta-analyzed depression subtypes with significantly correlated health traits. As the rG between fMDD and mMDD was very high in UKB (rG=0.93, P=4.83 × 10^−4^), this indicated that their additive genetic components were highly correlated, and that there was little justification to separate these groups on the basis of genetic stratification. The rG between fMDD and mMDD in the meta-analysis was out of bounds (rG=1.56) and non-calculable in GS:SFHS due to the near-zero h^2^_SNP_ of mMDD (as the h^2^_SNP_of both traits is included in the denominator term of the rG estimate)^36^. The lack of correlations surviving mMDD in LDSR analysis is therefore likely owing to reduced statistical power (μχ^2^=0.996) as a result of a smaller sample size. Figure 3 shows the genetic correlation of meta-analyzed depression with nominally and significantly correlated health traits.

**Figure 3.**
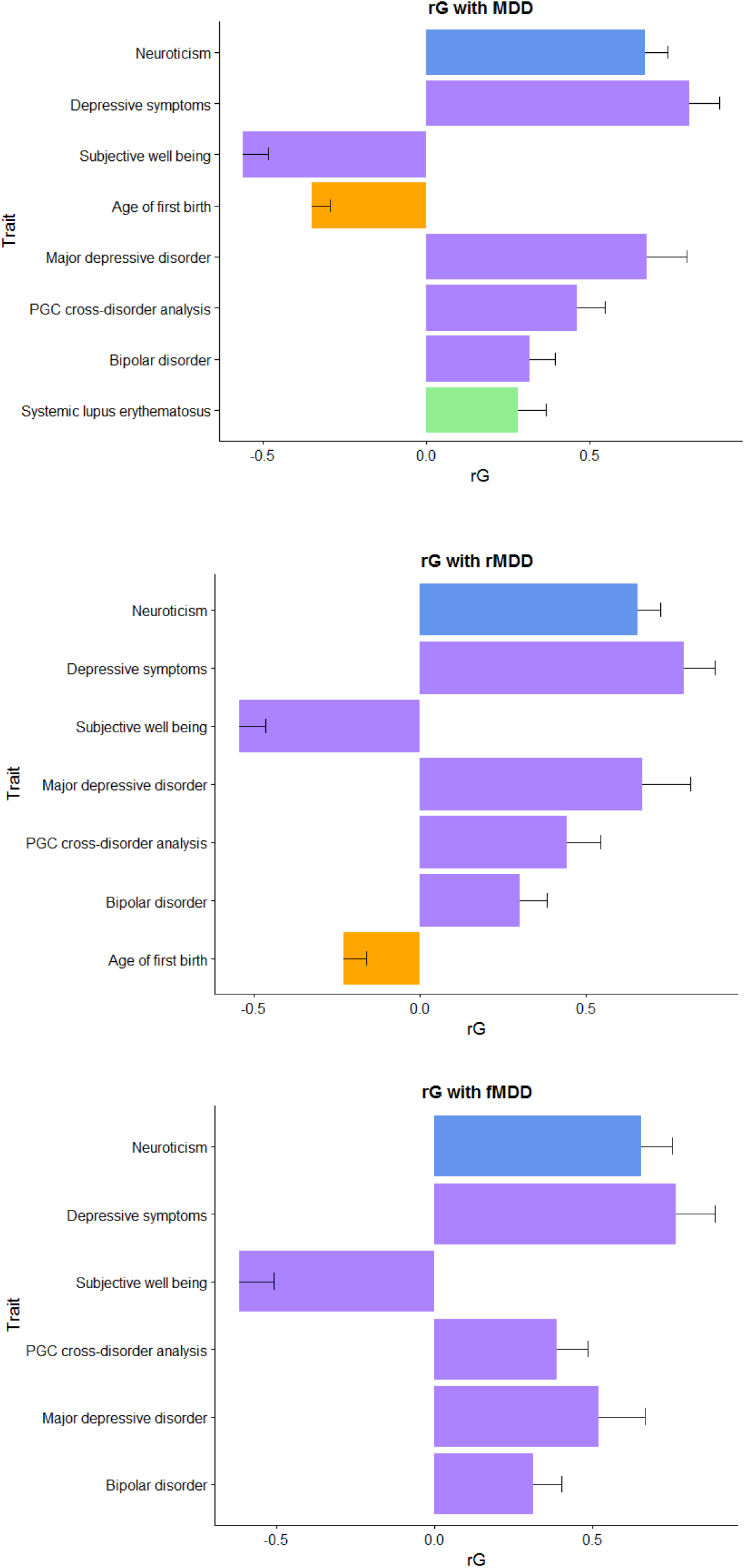
Genetic correlation (rG) between meta-analyzed MDD subsets and other health-related traits, derived using GWAS summary statistics and LD score regression. Traits presented showed a significant rG with MDD subsets after multiple testing correction (FDR p≤0.05) and are coloured by category (personality, psychiatric, reproductive and autoimmune). No rG between mMDD and other health-related traits survived multiple testing correction.

### Polygenic profiling analysis

The GWAS results from the MDD phenotype in 3 discovery samples (PGC MDD29, UKB and GS:SFHS) were used to build polygenic profile scores (PGS) in GS:SFHS and UKB, incorporating SNPs with a discovery sample association P-value cut-off of P_T_≤0.01, P_T_≤0.05, P_T_≤0.1, P_T_≤0.5, and all SNPs (P_T_≤1). Results from P_T_≤0.05 are shown in Figure 4, as this P_T_ explained the most variance in the target datasets. Results from all P_T_ are shown in Supplementary Tables 18-21. PGS derived using information from the PGC MDD29 GWAS yielded significant associations between depression phenotype and PGS across almost all thresholds in both GS:SFHS and UKB (with the exception of mMDD in GS:SFHS). PGS derived using information from the UKB MDD GWAS yielded significant associations with MDD, rMDD and mMDD phenotypes in GS:SFHS at all thresholds, however fMDD did not survive multiple testing correction at any P_T_. PGS derived using information from the GS:SFHS MDD GWAS yielded significant associations with MDD, rMDD and fMDD phenotypes in UKB at all thresholds except P_T_≤0.01, presumably due to the low number of SNPs contributing to the score, however mMDD did not survive multiple testing correction at any P_T_. Across all associations, the largest proportion of variance explained in GS:SFHS was 0.66% for fMDD using the MDD polygenic score derived using SNPs at P_T_≤0.5 using weights from the PGC MDD29 GWAS. The largest proportion of variance explained in UKB was 0.72% for fMDD, again using SNPs at P_T_≤0.5 using weights from the PGC MDD29 GWAS.

**Figure 4.**
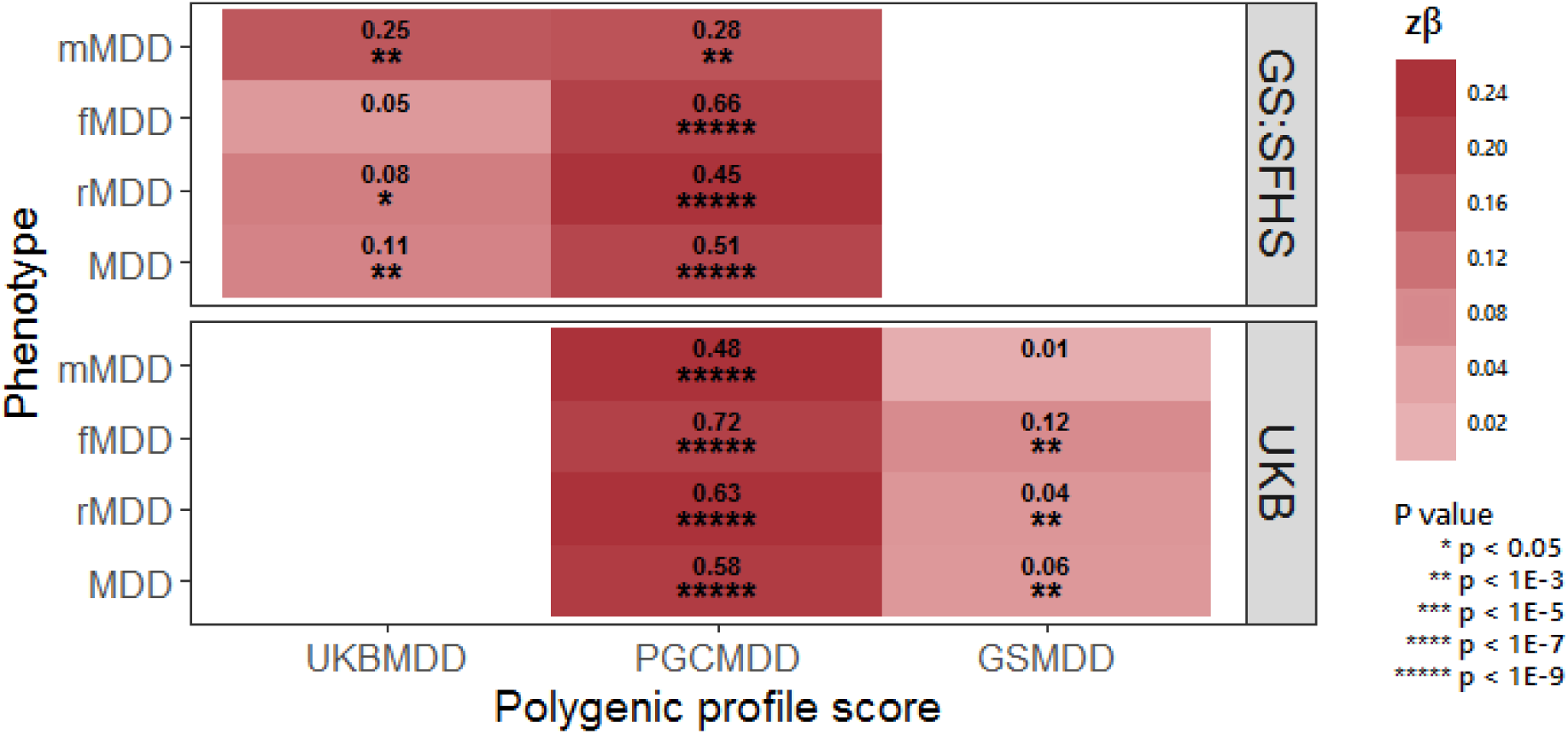
Heat map of associations between the polygenic profiles scores (PGS) for major depressive disorder (MDD), derived from Psychiatric Genomics Consortium (PGC) MDD29, UK Biobank (UKB) and Generation Scotland: The Scottish Family Health Study (GS:SFHS), and MDD subsets in UKB and GS:SFHS. Stronger associations are indicated by darker shades. The amount of variance (%) explained by PGS is indicated for each association. Further information can be found in Supplementary Tables 15-18.

## Discussion

For many years, the depressed phenotype has been refractory to genetic inquiry due to issues regarding statistical power. Recently, studies have successfully identified loci associated with depression by either refining the phenotype or by substantially increasing the sample size. In this study, we compared these strategies using data from two large UK cohorts (n_max_=43 062) to ascertain which approach may prove more advantageous for investigating the genetic architecture in each subgroup. This was achieved using techniques designed to interrogate complex traits, namely: meta-analysis, SNP-based heritability analysis, the genetic correlation with other health-related traits and variance explained by polygenic profile scores for depression.

A single genome-wide significant locus was identified in the meta-analysis of GS:SFHS and UKB for male depression. The locus at chromosome 3p22.3 includes *TMPPE, CRTAP* and *GLB1* genes; all three genes were significant in gene-based testing. *CRTAP* and *GLB1* are members of the CNTN1 PPI subnetwork, identified through gene set analysis. Whilst *CRTAP* and *GLB1* have not previously shown association with psychiatric disorders, the protein encoded for by *CNTN1* may play a role in the formation of axon connections in the developing nervous system^60^. Furthermore, the CNTN1 PPI subnetwork also contains *HTR1A*, which encodes for a serotonin (5-HT) receptor subtype that binds endogenous 5-HT^61^. Investigation of gene-expression databases suggests that the region on 3p22.3 contains eQTL for both *GLB1* and *CRTAP*. Akin to similarly sized published GWAS of depression^2,9^, we did not identify any genome-wide significant findings in our unstratified analysis, in individuals with recurrent depression or in the female-only analysis. This suggests that the phenotypic refinement demonstrated by the CONVERGE consortium which addressed several sources of heterogeneity – recurrence, sex and ancestral – has more utility than implementing recurrence and sex separately^6^.

In the SNP-based heritability analysis using GREML-based methods, no GS:SFHS MDD subset had a significant heritability estimate; UKB MDD subsets had significant heritability estimates although no subset showed increased heritability. Previous work exploring the heritability of MDD in GS:SFHS using Markov-Chain Monte-Carlos generalized linear mixed models (MCMCglmm) applied to pedigree data found that recurrent MDD and MDD in females had higher heritability point estimates, although confidence intervals still overlapped. However, both SNP-based estimates shown here and pedigree-based estimates from Fernandez-Pujals *et al^62^* are consistent with previous h^2^ estimates from the literature of estimate from Lee *et al*^63^ in PGC MDD (h^2^_SNP_ = 0.21, SE = 0.02) and Sullivan *et al*^64^ (h^2^ = 0.37, CI = 0.31 – 0.42).

We next investigated whether the subsets showed differing patterns of genetic correlations (rG) with other health-related traits. The bivariate correlation between unstratified MDD and health-related traits revealed eight statistically significant correlations. These included significant genetic correlations with a number of previously reported, well-established relationships such as neuroticism^65-67^, bipolar disorder^51,63^, PGC cross-disorder^63^, depressive symptoms^7,68^ and subjective well-being^69^. Relationships between MDD and both age at first birth and SLE have been previously reported although these have been based on phenotypic correlations^70-72^. Recurrent MDD and fMDD showed a very similar pattern of association (Supplementary Table 14) for personality and mood-related traits, and in both cases, the traits identified as being genetically correlated with r/fMDD were identical and in the same direction of association. In contrast to these results, bivariate genetic correlations between mMDD and health-related traits were all non-significant after adjustment for multiple testing.

Finally, we investigated the association of polygenic profile scores derived from three MDD GWAS with MDD subtypes, and how much trait variance these scores explained. The association between PGC-derived polygenic profile scores for MDD (derived after the elimination of common individuals) and MDD traits from GS:SFHS and UKB were significant for almost every unstratified and stratified depression phenotype, with the exception of MDD in males only in GS:SFHS, presumably due to this subset being much smaller in sample size (n=7 700). Scores derived from GWAS of MDD in UKB and GS:SFHS generally cross-associated in phenotypes from the alternate cohort (GS:SFHS MDD ~ UKB PGS; UKB MDD ~ GS:SFHS PGS), with the exception of fMDD in GS:SFHS and mMDD in UKB. In addition, PGS from GS:SFHS did not associate with any UKB MDD subtype at any P_T_, presumably due to extremely low statistical power in the GS:SFHS mMDD GWAS. Findings were generally more significant in UKB compared to GS:SFHS, and polygenic risk scores generated from the PGC MDD29 GWAS showed the strongest associations with all MDD subtypes. This likely reflects the increased size of these discovery and target GWAS datasets contributing to greater weight accuracy and increased statistical power to detect an association. However, the overlapping confidence intervals within GS:SFHS and UKB suggest that generally there were no detectable difference in the genetic architecture of these subsets.

There are several limitations to this study. Firstly, the sample size for MDD and rMDD groups are very similar (n=43 062 and n=39 556, respectively), therefore perhaps rMDD is not the best stratifier in this sample. The sample size of all depressed cases could have been increased by including individuals with mild depressive/manic symptoms (n=26 847), however as case classification was based on very few items (two symptoms and help-seeking behavior), it wasn’t possible to determine whether mild symptoms should be classified as cases or controls^17^, therefore including these individuals could introduce further phenotypic heterogeneity. The heterogeneity of the models used to adequately account for differential family and population structure in GS:SFHS and UKB is not ideal, however the genetic correlation of MDD GWAS summary statistics from the two samples was rG(SE) = 0.997(0.26), suggesting that perhaps this is not so problematic. The higher prevalence of females in GS:SFHS caused a gender imbalance in the sample sizes, resulting in lower statistical power for the males only analysis.

Overall our results suggest that there was little benefit to stratifying depression by either sex or recurrence for currently available data sizes. Extreme differences between sexes, such as opposite directions of effect in the two sexes, would have to exist to necessitate their analysis separately. The power implications of stratifying on these traits is likely to out-weigh the identification of such loci. In situations where the effect of a SNP is only found in one sex and zero effect in the other, such as rs4778037 in this study, it is still better to analyze sexes together to reduce the multiple testing burden of separate analyses. The increased trait variance explained demonstrated by using the largest available training and discovery datasets (PGC MDD29 and UKB MDD, respectively) in polygenic profiling supports increasing the sample size over phenotypic refinement. Similarly, the lack of discernible difference between h^2^_SNP_ and rG estimates between depression subtypes suggests that the best approach, currently, is to maximize the sample size in order to reduce sampling error and obtain more accurate point estimates. However large single population cohorts do have intrinsic merit, as demonstrated previously by CONVERGE^6^. Indeed, the relative ancestral homogeneity between GS:SFHS and UKB may have contributed to our identification of a genome-wide significant loci for depression in males. This finding will require further replication.

Phenotypic stratification still has plenty of scope for aiding the tractability of genetic analysis in depression. There are many traits which could be used to create subgroups including treatment response, overlap with other health-related traits and disease severity and perhaps these will be more successful than sex and recurrence. Recently, age-at onset^9^ and use of polygenic risk scores derived from health-related traits^73^ have been shown to result in subsets of depression with improved heritability. Our study describes the methods that can be applied to validate subsets for genetic analyses and concludes that stratification on the basis of sex and recurrence is not an optimal strategy however, it is worth noting that due to the limitations of statistical power with current sample sizes, the performance of the stratified phenotypes presented here is a lower bound of the stratification strategy.

## Acknowledgements

Generation Scotland received core funding from the Chief Scientist Office of the Scottish Government Health Directorate CZD/16/6 and the Scottish Funding Council HR03006. Genotyping of the GS:SFHS samples was carried out by the Genetics Core Laboratory at the Wellcome Trust Clinical Research Facility, Edinburgh, Scotland and was funded by the UK’s Medical Research Council and the Wellcome Trust (Wellcome Trust Strategic Award “STratifying Resilience and Depression Longitudinally” (STRADL) (Reference 104036/Z/14/Z). YZ acknowledges support from China Scholarship Council. IJD is supported by the Centre for Cognitive Ageing and Cognitive Epidemiology which is funded by the Medical Research Council and the Biotechnology and Biological Sciences Research Council (MR/K026992/1). AMMcI and T-KC acknowledges support from the Dr Mortimer and Theresa Sackler Foundation. This work has made use of the resources provided by the Edinburgh Compute and Data Facility (ECDF). (http://www.ecdf.ed.ac.uk/).

We are grateful to all the families who took part, the general practitioners and the Scottish School of Primary Care for their help in recruiting them, and the whole Generation Scotland team, which includes interviewers, computer and laboratory technicians, clerical workers, research scientists, volunteers, managers, receptionists, healthcare assistants and nurses. Ethics approval for the study was given by the NHS Tayside committee on research ethics (reference 05/S1401/8)

This research has been conducted using the UK Biobank Resource – application number 4844.

